# Tolerance of nonsynonymous variation is closely correlated between human and mouse orthologues

**DOI:** 10.1101/657981

**Authors:** George Powell, Michelle Simon, Sara Pulit, Ann-Marie Mallon, Cecilia M. Lindgren

**Affiliations:** Big Data Institute, Li Ka Shing Centre for Health Information and Discovery, University of Oxford; MRC Harwell Institute, Mammalian Genetics Unit, Oxfordshire, UK, OX11 0RD; Wellcome Centre for Human Genetics, University of Oxford, Oxford, UK; Medical and Population Genetics Program, Broad Institute of MIT and Harvard, Cambridge, MA, USA

## Abstract

Genic constraint describes how tolerant a gene is of nonsynonymous variation before it is removed from the population by negative selection. Here, we provide the first estimates of intraspecific constraint for mouse genes genome-wide, and show constraint is positively correlated between human and mouse orthologues (r = 0.806). We assess the relationships between mouse gene constraint and knockout phenotypes, showing gene constraint is positively associated with pleiotropy (ie an increased number of phenotype associations (R^2^ = 0.65)), in addition to an enrichment in lethal, developmental, and craniofacial knockout phenotypes amongst the most constrained genes. Finally, we show mouse constraint can be used to predict human genes associated with Mendelian disease, and is positively correlated with an increase in the number of known pathogenic variants in the human orthologue (R^2^ = 0.23). Our metrics of mouse and human constraint are available to inform future research using mouse models.

## INTRODUCTION

Pinpointing the genes, genetic variants, and biological pathways that underpin human disease remains a foremost focus of biomedical research today. Genome-sequencing has characterised human variation across global populations, and highlighted differences between genes with regard to the relative number of nonsynonymous variants they carry (Petrovski et al 2013; Lek et al 2016). This information has been used to estimate genic constraint, a description of how tolerant a protein-coding gene is to nonsynonymous variation before it is removed from the population by negative selection (Bartha et al 2018). Genes are more constrained if a) nonsynonymous variants have a high probability of affecting gene function, and b) there is strong purifying selection against the affect. Constrained genes are therefore characterised by a relative depletion of nonsynonymous variation. Multiple methods have been developed to quantify genic constraint in human populations (reviewed by Bartha et al 2018). The principle of each method is to quantify the difference between the relative number of nonsynonymous variants observed in each gene and either the genome-wide average (Petrovski et al 2013; Rackham et al 2015), or the expected number assuming neutral selection (Samocha et al 2014; Bartha et al 2015; Lek et al 2016; Fadista et al 2017; Cassa et al 2017). Constrained genes fall into a few known categories: some are essential for viability and development, while others associate with disease (Bartha et al 2018). Quantifying gene constraint can therefore help with the interpretation of personal genomes, including the identification of pathogenic variants.

Notably, genic constraint has not been estimated for mouse, which is the most widely utilised mammalian model organism for biomedical research (Rosenthal and Brown 2007; Justice and Dhillon 2016; Yue et al 2014), and as a result, the relationships in intraspecific constraint between human and mouse orthologues remains poorly understood. Quantifying differences in constraint between human and mouse orthologues could inform future clinical research using mouse models. This could be particularly pertinent for mouse humanization using CRISPR/Cas9 (Li et al 2014), and the clinical development of new drugs (Minikel et al 2019). Furthermore, quantifying mouse gene constraint would improve our understanding of the relationships between gene constraint and gene function in-vivo. The International Mouse Phenotyping Consortium (IMPC) is characterising mammalian gene function by systematically knocking out mouse genes and using a standardised pipeline to measure the resulting phenotypes across a spectrum of disease domains (Dickinson et al 2016; Smith and Epigg 2012; Karp et al 2015). This provides a unique resource to assess the global relationships between intraspecific gene constraint and gene function.

This study is the first to quantify intraspecific mouse gene constraint genome-wide and compare constraint between human and mouse orthologues, characterising genes that are most and least constrained in both species. We investigate the relationships between mouse gene constraint, mouse knockout phenotype, and human disease association of the human orthologue.

## RESULTS

### Identifying constrained genes in mice

Gene constraint is determined by a relative depletion of intraspecific nonsynonymous variation, and the power to detect constraint is therefore dependent on the number of variant sites within the population sample (Bartha et al 2018; Samocha et al 2014). We quantified constraint for mouse genes using whole genome sequences from the 36 laboratory mouse strains made publicly available by the Mouse Genomes Project (MGP) (Keane et al 2011). The number of variant sites between the MGP strains is sufficient to calculate constraint, and is comparable to the number of variant sites in human population samples (supplementary table 2). This is due to the phylogenetic distance between strains and the inbreeding of lineages which increases the probability of allele fixation by genetic drift (Adams et al 2015; Willoughby et al 2015).

We quantified constraint for 18,711 mouse genes as a functional Z-score (funZ). The premise of the funZ method is to quantify gene constraint by standardising the difference between the observed number of nonsynonymous (defined here as functional) variants in a gene and the expected number, predicted using a model trained on the number of synonymous (presumed non-functional and selectively neutral) variants. Genes with a higher funZ have relatively fewer functional variants than expected and are considered more constrained (figure 1). The funZ method is adapted from the missense Z-score method proposed by Samocha et al (2014) to make it suitable for application to the MGP dataset. There are two main methodological differences between the functional Z-score and the missense Z-score. First, we expand the definition of functional variation to include nonsense in addition to missense single nucleotide variants (SNVs). Second, we consider all variants in the MGP dataset that occur homozygous in one or more of the 36 mouse strains. Methodological differences result in variation between constraint metrics (Bartha et al 2018). We therefore calculated funZ for 17,367 human genes to standardise comparisons of constraint between human and mouse orthologues. We used the 1000 Genomes Project (1KGP) dataset (1000 Genomes Project Consortium 2015) as the source of variation to calculate human constraint to limit bias introduced by case control cohorts that are included in other publicly available datasets (Lek et al 2016). We consider all variants with a minor allele frequency (MAF) > 0.001 in the 1KGP dataset, thus increasing the probability that they occur homozygous within the population (Pemberton et al 2012; Allendorf 1986). FunZ is highly correlated with other metrics of human gene constraint (supplementary table 5).

**Figure 1.**
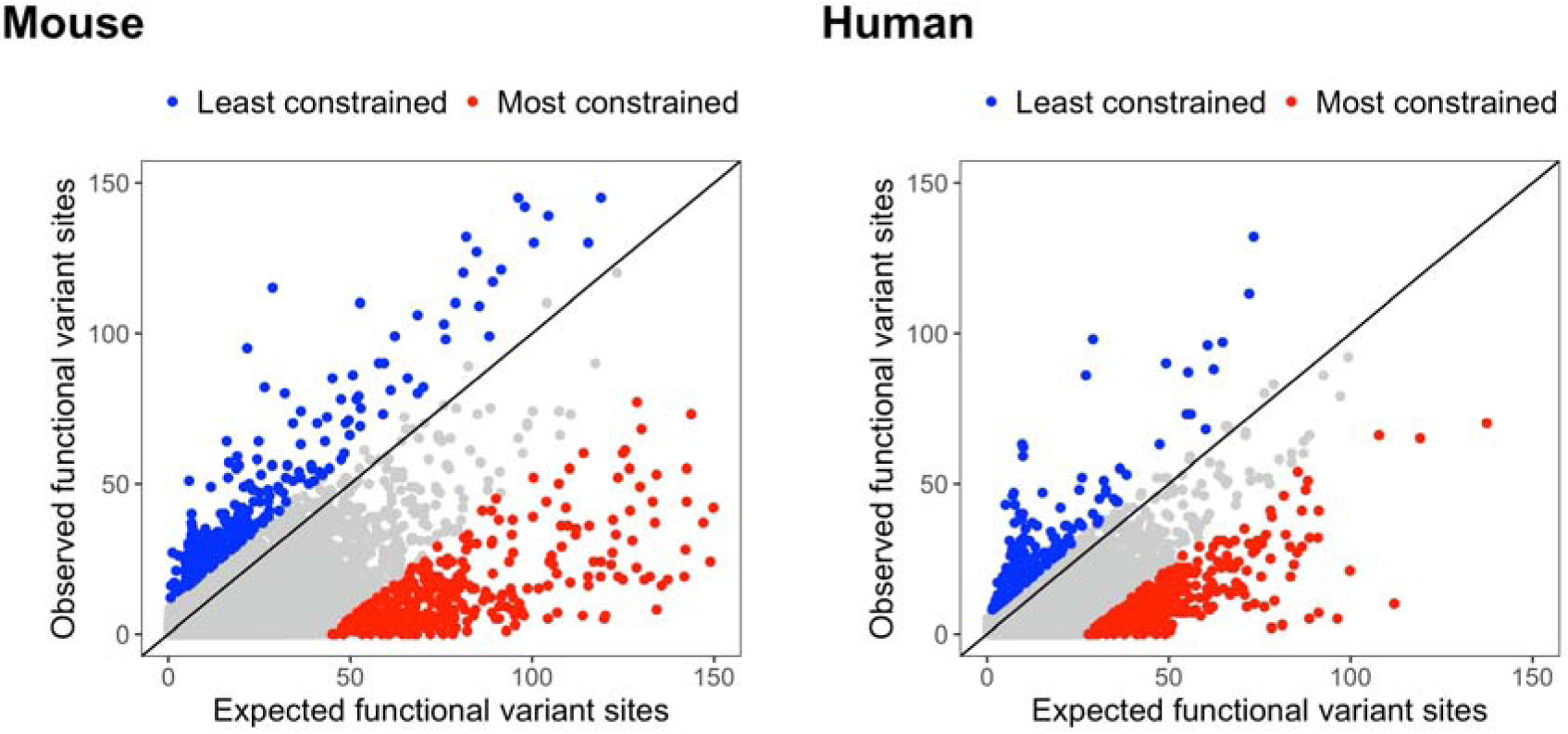
Scatter plots highlighting the relationship between the observed and expected common functional variant sites for mouse genes (n = 18,711) and human genes (n = 17,368). Variants are defined as “common” and “functional” if they have a MAF > 0.001, and are annotated as altering the amino-acid sequence of the protein. The expected number of functional variants is predicted with a model trained on the number of synonymous (presumed selectively neutral) variants. Constrained genes have proportionately fewer observed common functional variant sites than expected given no selection. The plots are annotated for the two percent most constrained and least constrained genes in red and blue respectively.

### Correlation in constraint between human and mouse orthologues

Orthologues are defined as genes for which speciation has occurred since divergence from the most recent common ancestor (Herrero et al 2016). They are classified as one-to-one when only one copy of the gene is found in each species; one-to-many when one gene in one species is orthologous to multiple genes in another species (ie the gene has multiplied in one lineage but not the other); or many-to-many when multiple orthologues are found in both species. We calculated constraint for 15,422 mouse, and 14,982 human orthologues defined by Ensembl (Zerbino et al 2018), using the MGP and 1KGP datasets respectively. Of these, 13,787 are defined as one-to-one orthologues, 902 human and 1,302 mouse genes are defined as one-to-many orthologues, and 293 human and 333 mouse genes are defined as many-to-many orthologues.

There is a significant positive correlation in constraint between human and mouse orthologues, computed as a Pearson’s product-moment correlation coefficient between funZ (r(16268) = 0.806, p<2.2e-16) (figure 2). This correlation is not, however, consistent between orthologous groupings as one-to-one orthologues are more closely correlated (r(13785) = 0.827, p<2.2e-16) than one-to-many (r(1477) = 0.536, p = 8.01e-111) and many-to-many orthologues (r(1002) = 0.148, p = 2.63e-06). We used Mann-Whitney U tests to assess differences in constraint between one-to-one, one-to-many, and many-to-many orthologues between human and mice. There is a significant difference in constraint between each group (p < 0.0001), with many-to-many orthologues the least constrained and one-to-one orthologues the most constrained (figure 3). This is consistent with previous work highlighting more constrained genes are less likely to have paralogues (Bartha et al 2015; Georgi et al 2013) and be copy number variable (Rudefer et al 2016).

**Figure 2.**
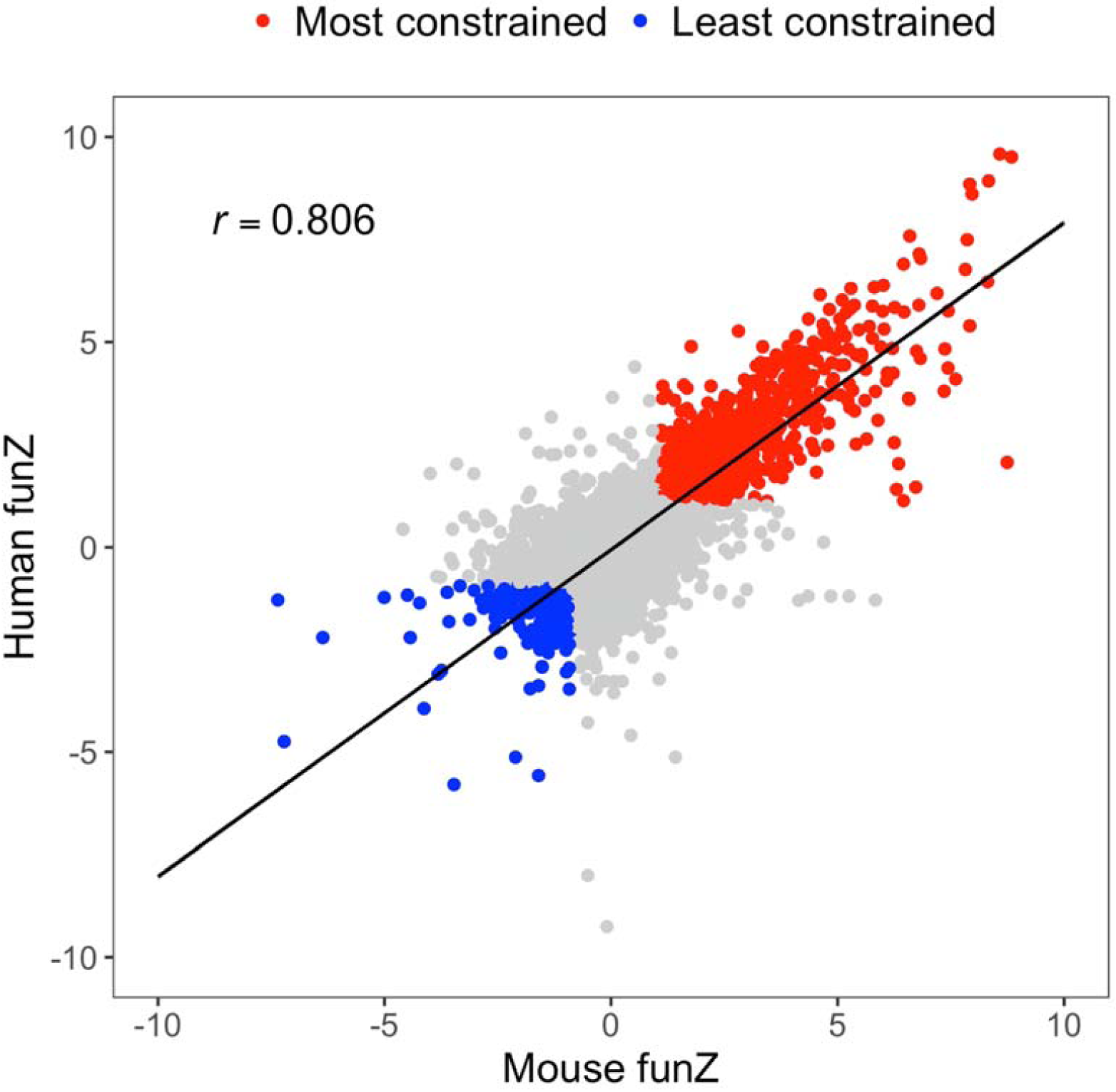
Constraint is correlated between human and mouse orthologues (r(16268) = 0.806, p<2.2e-16). Constraint was quantified for 15,422 mouse, and 14,982 human orthologues as funZ with a higher score indicating a greater degree of constraint. The most constrained orthologues in humans (n = 1,324) and mice (n = 1,321) were categorized as those that ranked among the top 10% for constraint in both species, and are annotated in red. The least constrained orthologues in humans (n = 327) and mice (n = 363) were categorized as those that ranked among the bottom 10% for constraint in both species, and are annotated in blue.

**Figure 3.**
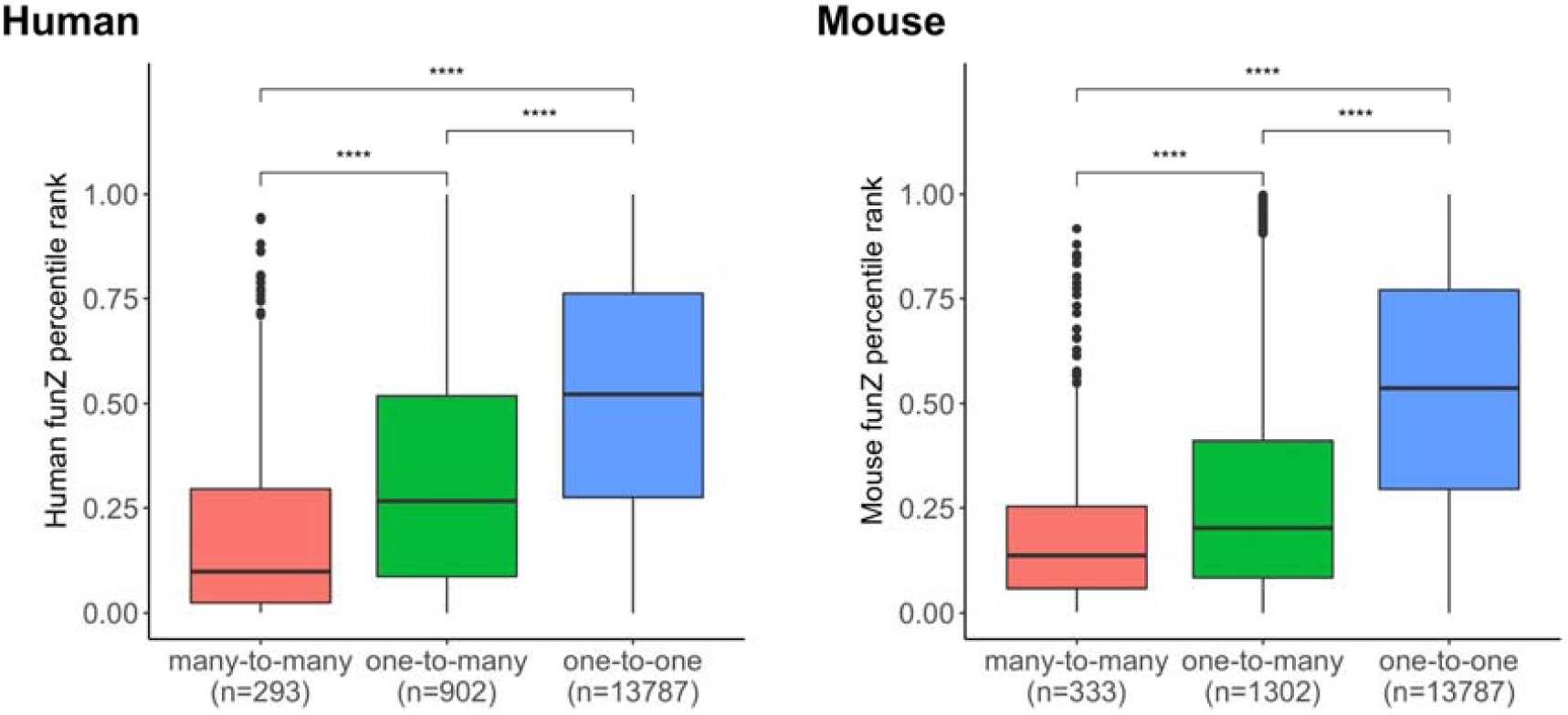
Distributions of constraint by orthology types (one-to-one, one-to-many, and many-to-many) for human and mouse orthologues. Constraint is quantified as funZ, with a higher score indicating a greater degree of constraint. Mann-Whitney U tests were used to assess differences between groups. There is a significant difference in constraint between each group (p < 0.0001), with many-to-many orthologues the least constrained and one-to-one orthologues the most constrained.

We assessed the relationship between intraspecific constraint (measured as funZ) and interspecific conservation (measured as the percentage of amino-acid sequence that matches between orthologous genes) by computing the Spearman’s Rank correlation. There is a significant positive correlation between mouse constraint and human-mouse conservation (n=16,270, rs=0.566, p<2.2e-16), and between human constraint and human-mouse conservation (n=16,270, rs=0.497, p<2.2e-16) (supplementary figure 2). This highlights that constrained genes are more likely to be conserved over evolutionary time (Bartha et al 2018).

### Gene constraint and knockout phenotype

We characterised the relationships between gene constraint and gene function by considering Mouse Phenotype ontology (MP) annotations from gene knockouts conducted by the IMPC (release 9.1). We grouped 5,486 gene knockouts studied by the IMPC by their top-level MP terms. Each gene was included a maximum of once for each top-level term grouping, and top-level terms with less than 50 associated genes were removed from the analysis. IMPC knockouts are subject to a standardised phenotyping pipeline; however, there is some variation in which phenotyping tests are performed due to differences in knockout lethality and funding limitations. We therefore compared funZ between all knockouts annotated with a top-level MP (i.e. knockouts that passed a significance threshold of 0.0001 in one of the associated phenotyping tests), with all knockouts that do not have the top-level MP annotation but were subject to one or more of the associated phenotyping tests. We used Mann-Whitney U tests with a Bonferroni correction for multiple testing to assess differences between the groups (figure 4, supplementary table 6). Eleven of the 21 top-level MP terms comprised genes that were significantly (p < 0.05) more constrained than genes that were tested for but did not have the top-level MP annotation, with the greatest difference for mortality/aging, craniofacial, and growth/size/body category phenotypes (figure 4, supplementary table 6). It is of note that the subset of 1,339 knockouts with no IMPC MP annotations are significantly less constrained than the average for all knockouts (p = 2.4e-16).

**Figure 4.**
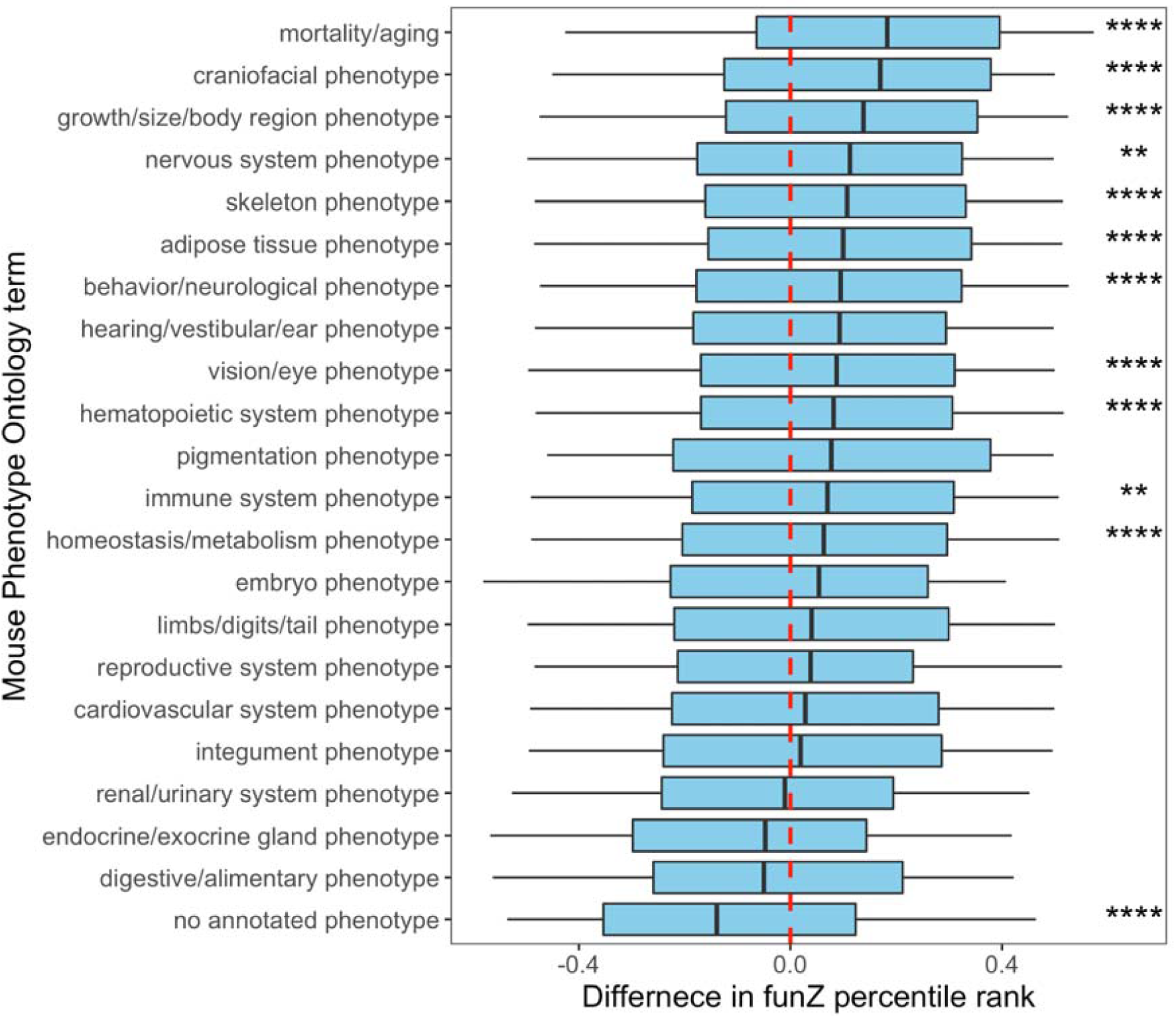
Differences in constraint between mouse genes associated with 21 top-level phenotype terms from the Mammalian Phenotype (MP) Ontology, and for knockouts with no annotated MP terms from the International Mouse Phenotyping Consortium (IMPC). We assessed 5,486 knockouts conducted by the IMPC. Constraint was quantified for each knockout as the percentile rank of funZ, with a higher score indicating a greater degree of constraint. The difference in funZ from each MP grouping was standardised against the median funZ of knockouts that have had one or more MP associated phenotyping test in the IMPC pipeline but are not annotated with the MP, indicated by the red line. Mann-Whitney U tests were conducted with a Bonferroni correction for multiple testing to assess significance between groups (* signify significance thresholds of 0.05, 0.01, 0.001, and 0.0001).

Genes can affect multiple, often seemingly unrelated, phenotypes, and this phenomenon is known as pleiotropy. We hypothesised that the more phenotypes a gene affects (the more pleiotropic a gene is), the more likely it is to be under selective constraint. To test this hypothesis, we assessed the relationship between gene constraint and the proportion of MP ontology annotations associated with the IMPC knockout for 5,486 genes. The proportion of MP ontology annotations associated with each knockout was calculated by dividing the total MP terms associated with each knockout by the potential number of MP terms (determined by the phenotyping tests that were performed). We binned knockouts by funZ from 1 to 100 with the least constrained genes in the 1st bin and the most constrained genes in the 100^th^ bin. We performed simple linear regression to predict the median proportion of MP terms per mouse knockout as a function of funZ percentile bin (figure 5). A significant regression equation was found (F(1, 98) = 140.5, p=1.2e-20) with an R2 of 0.59. The predicted proportion of MP terms is equal to 1.4e-02 + 2.3e-04 for each percentile increase in funZ. To assess whether this relationship is consistent for distantly related phenotypes we also performed simple linear regression to predict the median proportion of top-level MP terms per mouse knockout as a function of funZ percentile bin (figure 5). The MP ontology is a directed acyclic graph, and it is possible for one MP term to have multiple top-level terms (Eppig et al 2015). We therefore ensured only one top-level term was counted per MP term. A significant regression equation was found (F(1, 98) = 178.4, p=8.6e-24) with an R2 of 0.65. The predicted number of top-level MP terms is equal to 6.4e-02 + 9.4e-04 for each percentile increase in funZ. Our results highlight genic constraint is positively correlated with an increase in knockout phenotypes, indicating constrained genes are more likely to be pleiotropic.

**Figure 5.**
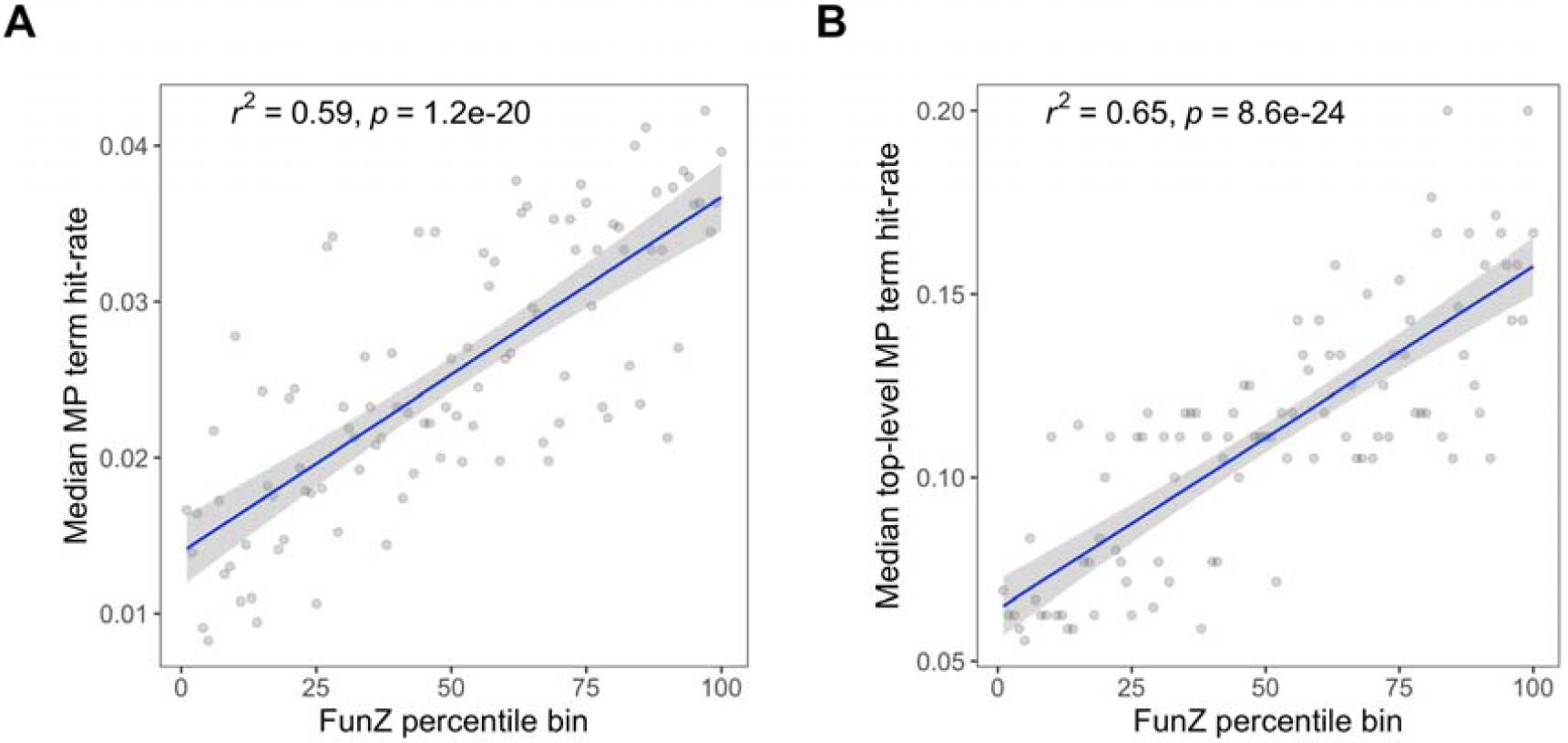
Mouse gene constraint is positively correlated with an increased number of knockout phenotypes. Mouse Phenotype Ontology (MP) terms associated with 5,486 mouse knockouts were obtained from the International Mouse Phenotyping Consortium. The MP term hit-rate for each knockout was calculated by adjusting the total MP terms associated with each knockout by the potential number of MP terms (determined by the phenotyping tests that were performed). Knockouts were binned from 1 to 100 based on their funZ percentile, with the least constrained genes in the 1^st^ bin and the most constrained genes in the 100^th^ bin. Regression lines are for (A) the median MP term hit-rate per knockout as a function of funZ percentile bin, and (B) the median top-level MP term hit-rate per knockout as a function of funZ percentile bin.

### Human disease association

Human gene constraint is positively correlated with disease association (Bartha et al 2018). We considered two hypotheses for assessing the relationships between mouse gene constraint and disease association of the human orthologue: 1) mouse gene constraint can be used to predict human orthologues associated with Mendelian disease; 2) mouse gene constraint is positively correlated with an increase in known pathogenic variants in the human orthologue.

We considered five lists of human genes associated with Mendelian disease to assess whether mouse gene constraint can be used to predict association to Mendelian disease of the human orthologue. The gene lists were curated by Petrovski et al (2013) using keyword searches in the Online Mendelian Inheritance in Man (OMIM) database, and have been used to assess the predictive performance of other constraint metrics including the RVIS (Petrovski et al 2013) and missense Z-score (Samocha et al 2014). Keyword searches included “haploinsufficiency”, “dominant-negative”, “de novo”, and “recessive”, in addition to a list of all OMIM disease genes. We used logistic regression to assess the difference in funZ between mouse one-to-one orthologues of human genes with no OMIM disease gene association (n=9,906), and each of the OMIM gene lists, and assessed predictive power as ROC (table 1). We benchmarked the predictive power of mouse funZ against funZ for the human gene, RVIS, missense Z-score, and pLI (table 1). Genes in each of the OMIM lists are significantly more constrained (measured as funZ, RVIS, missense Z-score and pLI) than genes with no OMIM disease gene association (table 1). Mouse orthologues of genes in each of the OMIM lists have a significantly higher funZ (are more constrained) than mouse orthologues of human genes with no OMIM disease gene association (figure 6, table 1). Mouse funZ has a similar predictive power to human funZ (table 1), with the difference in constraint most pronounced for the “haploinsufficiency” and the “de novo” gene lists (figure 6).

**Table 1.** Efficacy of funZ, RVIS, missense Z-score, and pLI in predicting gene lists from the Online Mendelian Inheritance in Man (OMIM) database. FunZ is calculated for the gene and the mouse orthologue. Keyword searches include “haploinsufficiency” (n=151), “dominant-negative” (n=317), “de novo” (n=383), and “recessive” (n=687), in addition to a list of all OMIM disease genes (n=1,917). We used logistic regression to assess the difference in constraint between genes with no OMIM disease gene association (n=9,906), and each of the OMIM gene lists.

| OMIM search term (n) |  | Missense Z-score | pLI | RVIS | Human funZ | Mouse funZ |
| --- | --- | --- | --- | --- | --- | --- |
| “haploinsufficiency”<br>(n=151), | Estimate<br>(Std error) | 3.40 (0.36) | 2.35<br>(0.22) | -3.23<br>(0.35) | 3.61<br>(0.37) | 3.47<br>(0.36) |
|  | P | 8.1e-21 | 1.5e-25 | 4.8e-20 | 1.5e-22 | 1.0e-21 |
|  | ROC | 0.74 | 0.76 | 0.73 | 0.75 | 0.75 |
| “dominant-negative”<br>(n=317) | Estimate<br>(Std error) | 1.73 (0.21) | 1.01<br>(0.13) | -1.60<br>(0.21) | 1.75<br>(0.21) | 1.50<br>(0.21) |
|  | P | 3.9e-16 | 8.9e-15 | 2.5e-14 | 9.8e-17 | 4.6e-13 |
|  | ROC | 0.64 | 0.62 | 0.63 | 0.64 | 0.62 |
| “de novo” (n=383) | Estimate<br>(Std error) | 2.24 (0.20) | 1.26<br>(0.12) | -1.55<br>(0.19) | 2.10<br>(0.20) | 2.23<br>(0.20) |
|  | P | 2.0e-28 | 1.0e-25 | 3.4e-16 | 3.3e-26 | 9.4e-29 |
|  | ROC | 0.67 | 0.65 | 0.62 | 0.66 | 0.67 |
| “recessive” (n=687) | Estimate<br>(Std<br>error) | -0.37 (0.14) | -0.39<br>(0.10) | -0.38<br>(0.14) | 0.68<br>(0.14) | 0.92<br>(0.14) |
|  | P | 6.9e-03 | 1.1e-04 | 5.8e-03 | 7.1e-07 | 4.6e-11 |
|  | ROC | 0.53 | 0.56 | 0.53 | 0.56 | 0.58 |
| all OMIM disease genes<br>(n=1,917). | Estimate<br>(Std<br>error) | 0.27 (0.09) | 0.04<br>(0.06) | -0.68<br>(0.09) | 0.87<br>(0.09) | 0.96<br>(0.09) |
|  | P | 1.3e-03 | 4.7e-01 | 5.4e-15 | 2.1e-23 | 1.2e-27 |
|  | ROC | 0.52 | 0.51 | 0.56 | 0.57 | 0.58 |

**Figure 6.**
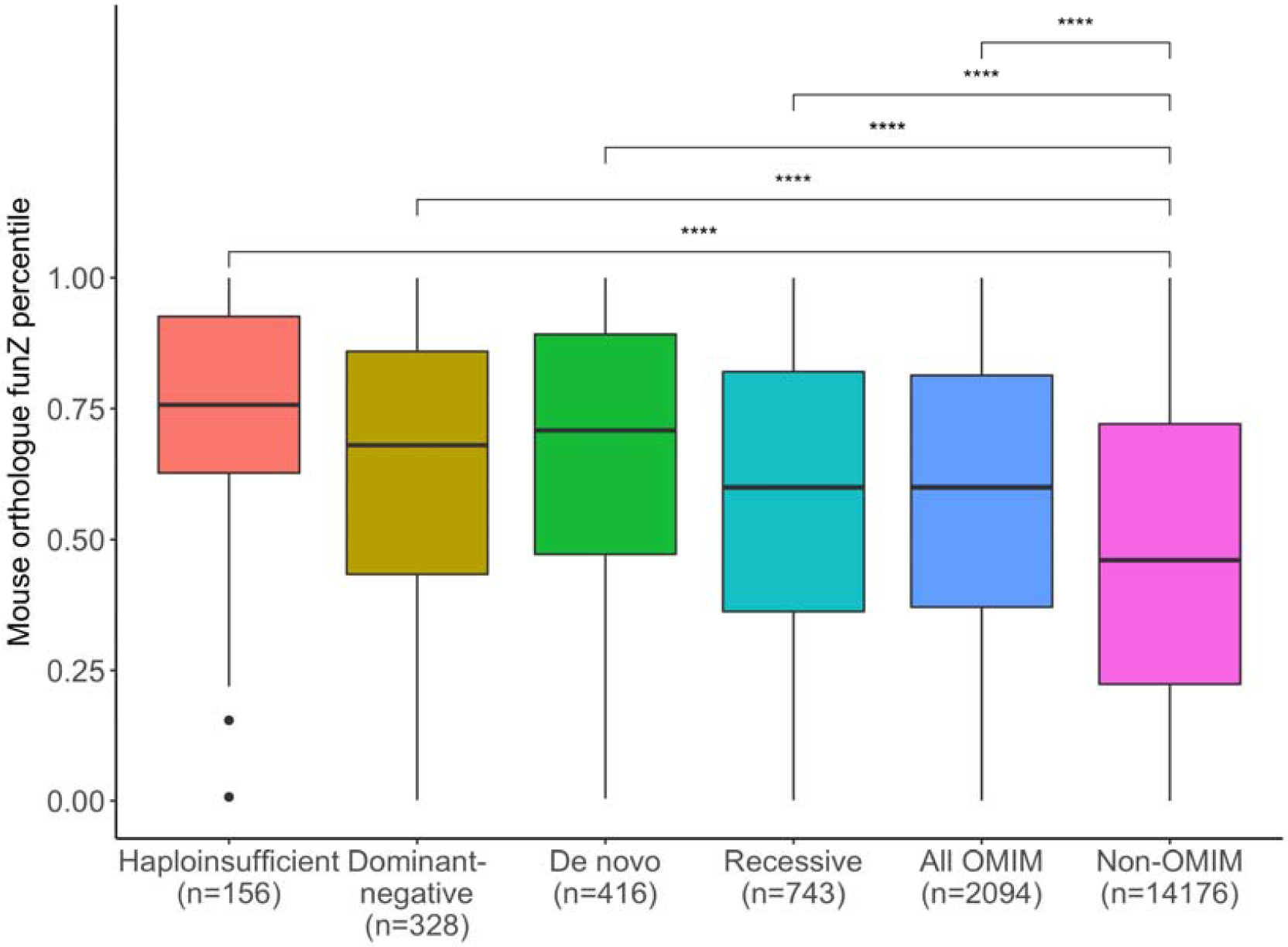
Distributions of constraint percentile for one-to-one mouse orthologues of human genes associated with Mendelian disease. Constraint was quantified for each gene as a percentile rank of funZ with a higher score indicating a greater degree of constraint. Mendelian disease gene lists were curated by Petrovski et al (2013) using key-word searches in the Online Mendelian Inheritance in Man (OMIM) database. Logistic regression models were used to assess the difference in constraint between each group and orthologues not included in any of the gene lists (non-OMIM). Mouse orthologues of human genes associated with Mendelian disease are significantly more constrained (p<0.0001) than mouse orthologues of human genes not include in any of the gene lists.

We assessed the relationship between mouse gene constraint and the number of known pathogenic variants in the human orthologue by considering 52,174 pathogenic variants from the ClinVar database (Landrum et al 2018). Human-mouse orthologues were binned from 1 to 100 based on their funZ percentile, with the least constrained genes in the 1^st^ bin and the most constrained genes in the 100^th^ bin. To account for differences in gene length, we averaged pathogenic variants in each gene per kb. We fit a simple linear regression to predict the mean number of pathogenic variants per kb as a function of funZ percentile bin for 15,680 mouse and 15,562 human orthologues (figure 7). A significant regression equation was found for mouse funZ (F(1, 98) = 28.64, p=5.7e-07) with an R2 of 0.226. The predicted number of pathogenic variants per kb = to 0.99 + 0.01 for each percentile increase in funZ in the mouse orthologue. This suggests gene constraint can be in part explained by variants in more constrained genes having an increased likelihood of being pathogenic.

**Figure 7.**
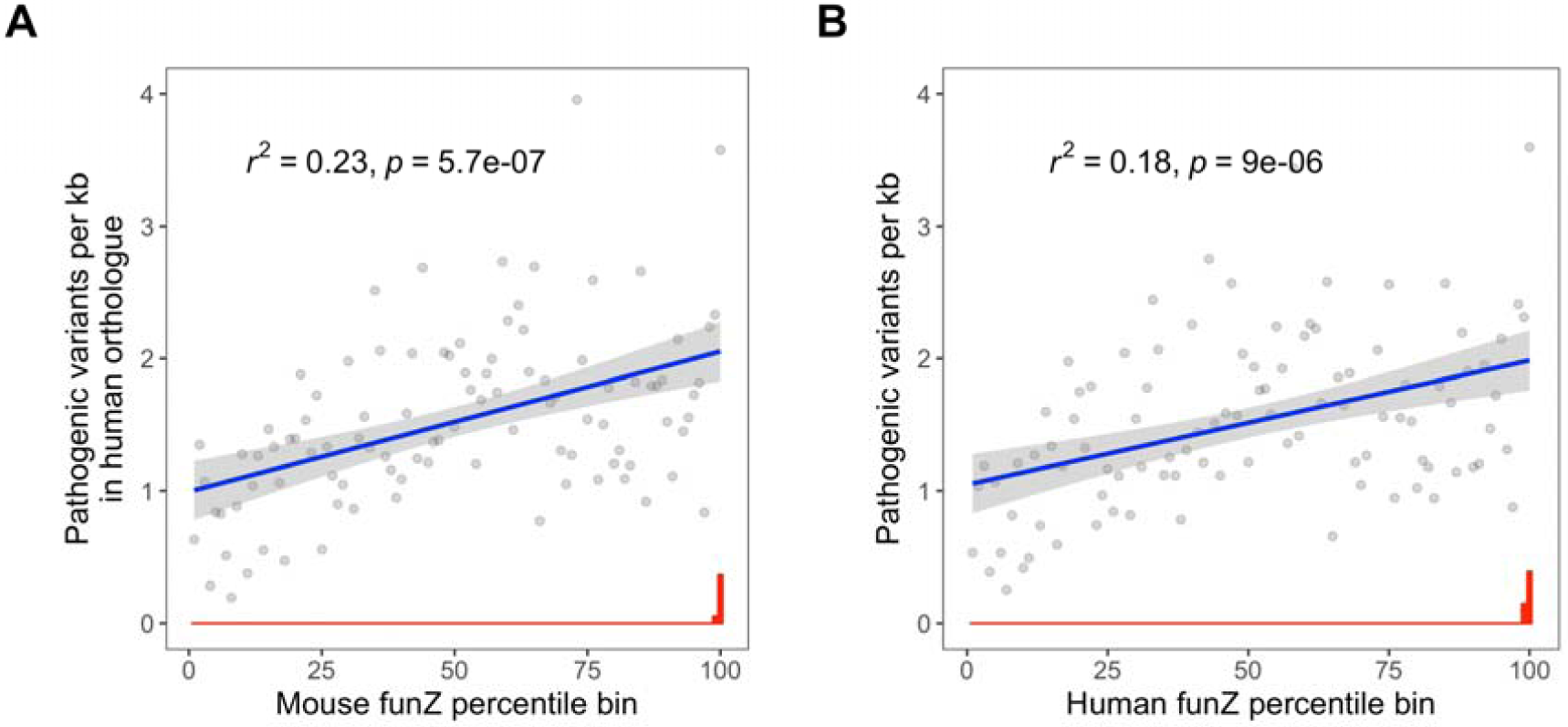
Mouse constraint is correlated with the number of known pathogenic variants in their human orthologues. Pathogenic variants were obtained from the ClinVar database (n = 52,174). 15,680 mouse and 15,562 human orthologues were binned from 1 to 100 based on their funZ percentile, with the least constrained genes in the 1st bin and the most constrained genes in the 100th bin. Regression lines are for A) the mean number of pathogenic variants per kb in the human orthologue as a function of mouse funZ percentile bin (p=5.7.e-07), and B) the mean number of pathogenic variants per kb as a function of human funZ percentile bin (p=9.1e-06). Standard error is highlighted in grey. The median number of pathogenic variants per kb for each percentile bin is given in red, and highlights an enrichment of known pathogenic variants in the two percentiles containing the most constrained genes in humans and mice. Gene constraint can be in part explained by variants in more constrained genes having an increased likelihood of being pathogenic.

## DISCUSSION

We quantified mouse gene constraint genome-wide, and compared intraspecific constraint between human and mouse orthologues. Our research has three main findings: First, genic constraint is positively correlated between human and mouse orthologues. This correlation is not, however, consistent between orthology types. We show that constraint is more closely correlated between one-to-one orthologues than one-to-many and many-to-many orthologues. This is consistent with previous work highlighting more constrained genes are less likely to have paralogues (Bartha et al 2015; Georgi et al 2013) and be copy number variable (Rudefer et al 2016). Second, mouse gene constraint is positively correlated with an increased number of knockout phenotype annotations, suggesting genes that are pleiotropic (ie influence multiple phenotypes and pathways) are more likely to be under selective constraint. We furthermore highlight an enrichment of constrained genes in mice that are associated with lethality, developmental and craniofacial knockout phenotypes. Third, mouse constraint can be used to predict human genes associated with Mendelian disease, and is positively correlated with an increase in the number of known pathogenic variants in the human orthologue. This is best explained by the correlation in constraint between mouse and human orthologues, as human gene constraint has been previously shown to correlate with disease association and pathogenic variant enrichment (Bartha et al 2018).

Estimates of gene constraints are dependent on methodological assumptions and the source of genetic variation on which they are based (Bartha et al 2018). To calculate constraint for mouse genes we used sequence variation from 36 mouse strains that have been inbred to achieve homology of genetic backgrounds (Adams et al 2015). In diploid organisms, selection strength, and therefore constraint, is influenced by penetrance and zygosity (Fuller et al 2018). For example, variants may be under stronger negative selection in homozygous individuals than heterozygous if there is lower penetrance associated with heterozygosity. Inbreeding increases homozygosity and the probability that deleterious recessive alleles will be removed from the population by negative selection (Willoughby et al 2015). Our estimate of mouse gene constraint is therefore biased towards identifying genes that are intolerant of homozygous variation. To account for this in our estimate of human gene constraint we only considered variants with an MAF > 0.001, thus increasing the probability that they occur homozygous within the population (Pemberton et al 2012; Allendorf 1986).

We observed a greater correlation in intraspecific constraint between human and mouse orthologues compared with the correlation between intraspecific constraint and interspecific conservation. This has two potential explanations: First, selection pressure and therefore constraint can change over evolutionary time, and this may have led to deviation in the amino-acid sequences of orthologous genes since the lineages diverged. Second, there is regional variability in constraint within genes due to differences in the functional importance of loci (Havrilla et al 2018). This could result in within-gene deviation in the amino acid sequence at loci that are of less functional importance.

In conclusion, the positive correlation in constraint between human and mouse orthologues indicates a positive correlation in functional importance between orthologous genes. The strength of this correlation supports the use of mouse as a model for understanding the mechanistic basis of gene function and human monogenic disease.

## METHODS

### Defining genes, quality variants, and coding consequences

We used two highly curated publicly available datasets as sources of genetic variation to calculate constraint for human and mouse genes: the Mouse Genomes Project dataset for mice (Keane et al 2011), and the 1000 Genomes Project (Phase 3) dataset for humans (1000 Genomes Project Consortium 2015). We considered all protein-coding genes with a HUGO Gene Nomenclature Committee name, and defined the coding sequence for each gene by their Ensembl canonical transcript (release 94) (Zerbino et al 2018). We considered all single nucleotide variants (SNVs) with “PASS” filter status as described by the 1000 Genomes Project and Mouse Genomes Project (1000 Genomes Project Consortium 2015; Keane et al 2011). Genes were filtered that do not have one or more SNV in their canonical transcript. Measurements of constraint are biased towards longer genes with more variants, and we therefore removed genes with a canonical transcript > 1.5 kb, or more than 300 SNVs. This left 17,367 human and 18,710 mouse genes for analysis. Orthologous genes were defined by Ensembl (release 94) (Zerbino et al 2018). The final dataset consists of 14,982 human and 15,422 mouse genes with one or more orthologue, including 13,787 one-to-one orthologues; 1,479 one-to-many orthologues consisting of 902 unique human and 1302 unique mouse genes respectively; and 1,004 many-to-many orthologues consisting of 293 unique human and 333 unique mouse genes respectively.

We classified SNVs as “functional” and “nonfunctional” based on their annotated consequences for the amino-acid sequence (supplementary table 1). Functional variants are assumed to change the amino-acid sequence, and non-functional variants are assumed to be silent. The coding consequences of SNVs in the 1000 Genomes Project and Mouse Genomes Project datasets were determined using the Ensembl Variant Effect Predictor (v94.5) (McLaren et al 2016). One consequence was determined per SNV using the “--pick” argument which prioritises annotations by canonical transcript status. We defined missense and nonsense variants as functional, and synonymous variants as non-functional.

### Calculating sequence-specific probabilities of variation

The probability of a DNA sequence incurring a substitution mutation is in part dependent on its local sequence context (Aggarwala and Voight 2016). Consistent with the missense Z-score method (Samocha et al 2014), we considered the trinucleotide context for calculating gene-specific probabilities of substitution (ie the probability of Y_1_ in the trinucleotide XY_1_Z mutating to Y_2_ in the trinucleotide XY_2_Z is dependent on X and Z). We estimated the 192 relative substitution rate probabilities of the middle base in each of the 64 potential trinucleotides for humans and mice by considering the intergenic SNVs in the 1000 Genomes Project and Mouse genomes Project datasets, and using human to chimpanzee (*Pan troglodytes*) and mouse (*Mus musculus*) to *Mus Caroli* alignments from Ensembl (release 94) to infer the mutational direction for each SNV (ie which of the reference and alternate bases is the “ancestral” and “mutant”). We inferred the ancestral and mutant bases for each SNV following two assumptions: a) the ancestral base is the reference base, or the alternate base if the alternate base is shared with the related species; b) the mutant base is the alternate base or the reference base if the alternate is shared with the related species. For each trinucleotide, we calculated the relative probabilities of substitution by dividing the observed number of intergenic trinucleotide changes by the number of the trinucleotide in the intergenic ancestral sequence. Trinucleotide mutation rate probabilities estimated for the human and mouse lineages are highly correlated (r(190)=0.995, p=2.0e-192)(supplementary figure 1). We used the trinucleotide mutation rate probability tables to estimate the probabilities of incurring synonymous and functional mutations for human and mouse genes by considering the coding consequences for each potential substitution in the canonical transcript, and totalling the trinucleotide specific probabilities of mutation. Trinucleotide mutation rate tables and gene-specific probabilities of mutation for humans and mice are provided in the supplementary information.

### Calculating regional and intron mutation rates

Mutation rates vary throughout the genome (Hodgkinson and Eyre-Walker 2011). We therefore estimated the regional mutation rate for each gene by counting the number of SNVs within the genes start and end coordinates plus 1Mbp upstream and downstream, and dividing by the difference between the start and end coordinates plus 2,000,000. In addition, we estimated the intron mutation rate for each gene canonical transcript by dividing the number of intron SNVs (MAF > 0.001) with the sum of intron lengths. Regional mutation rates for human and mouse genes are provided in the supplementary information.

### Calculating gene constraint as the functional Z-score (funZ)

We quantified constraint for mouse and human genes as a functional Z-score. FunZ is calculated in a two-stage process. First, we built a model to predict the number of SNVs in each gene assuming no selection pressure by regressing the number of common (MAF > 0.001) synonymous variants against the genes sequence-specific probability of synonymous mutation, regional mutation rate, and intron mutation rate. Model fit and covariate significance are provided in supplementary table 3. To compare the impact of MAF on the results, we calculated constraint for human genes across a range of MAF thresholds (MAF > 0.001, MAF > 0.0005, and MAF > 0.0001), and funZ is closely correlated between the results (supplementary figure 4). The Mouse Genomes Project dataset has a greater ratio of synonymous variants to functional variants compared to the 1000 Genomes Project Dataset. This can be explained by the increased probability of synonymous fixation by genetic drift during the selective breeding of inbred strains (Willoughby et al 2015). To account for this we divided the number of synonymous variants in each gene in the Mouse Genomes Project dataset by two before regression. Second, we use this model to predict the expected number of functional SNVs in each gene, given neutral selection, by substituting in the genes sequence-specific probability of functional mutation. We standardised the difference between the observed and expected number of common functional variants for each gene as a Z-score (funZ). Genes with a higher funZ have relatively fewer common functional variants than expected and are considered more constrained (figure 1).

### Correlation between human and mouse orthologues, and with other measures of intraspecific constraint and interspecific conservation

Human-mouse orthogues were defined by Ensembl (release 94) (Zerbino et al 2018), and correlation in constraint between orthologous genes was calculated as a Pearson’s product-moment correlation coefficient between funZ. We calculated the Spearman’s Rank correlation between human constraint measured as funZ, and previously published measures of intraspecific constraint (RVIS, missense Z-score, and pLI) (supplementary table 5). We calculated the Spearman’s Rank correlation between human and mouse constraint measured as funZ, and interspecific conservation measured as the mean percentage of amino-acid sequence that matches between orthologues (Query % ID and Target % ID) (Zerbino et al 2018) (supplementary figure 2).

### Assessing mouse constraint and knockout phenotype

We investigated the relationship between gene constraint measured as funZ and gene function by considering knockout phenotypes for 5,486 genes from the IMPC)(release 9.1. Knock-out phenotypes are quantified using a standardised pipeline and annotated in the MP(Smith and Epigg 2012). We grouped genes by associated top-level MP term. Each gene was included a maximum of once in each group. We discarded top-level MP terms with less than 50 associated knockouts. We also curated a list of genes that have been knocked out by the IMPC, but have no MP annotations. IMPC knockouts are subject to different phenotyping pipelines due to due to differences in lethality and ethical limitations. We therefore compared funZ between all knockouts annotated with a top-level MP (ie knockouts that passed a significance threshold of 0.0001 in one of the associated phenotyping tests), with all knockouts that do not have the top-level MP annotation but were subject to one or more of the associated phenotyping tests. We used Mann-Whitney U tests with a Bonferroni correction for multiple testing to assess differences between groups.

We investigated the relationship between gene constraint and the proportion of MP terms associated with the IMPC knockout, to serve as a proxy for the pleiotropic effect of a gene. The proportion of MP ontology annotations associated with each knockout was calculated by dividing the total MP terms associated with each knockout by the potential number of MP terms (determined by the phenotyping tests that were performed). We binned 5,486 IMPC knockouts by funZ from 1 to 100 with the least constrained genes in the 1^st^ bin and the most constrained genes in the 100^th^ bin. We performed two simple linear regressions: 1) to predict the median proportion of unique MP terms per mouse knockout as a function of funZ percentile bin, and 2) to predict the median proportion of unique top-level MP terms per mouse knockout as a function of funZ percentile bin. The MP ontology is a directed acyclic graph, and it is possible for one MP term to have multiple top-level terms. We therefore ensured only one top-level MP term was counted per MP term.

### Assessing mouse constraint and human disease gene association

We benchmarked the ability of human and mouse funZ to predict genes associated with Mendelian disease against the publicly available constraint metrics RVIS (Pertovski et al 2013), missense Z-score (Samocha et al 2014), and pLI (Lek et al 2016). We considered five lists of human genes associated with human disease curated by Petrovski et al (2013) using keyword searches in the Online Mendelian Inheritance in Man database. Keyword searches included “haploinsufficiency”, “dominant-negative”, “de novo”, and “recessive”, in addition to a list of all OMIM disease genes. We used univariate logistic regression models to assess the difference in constraint measured as funZ, RVIS, missense Z-score, pLI between genes with no OMIM disease gene association, and each of the OMIM gene lists, in addition to a multivariate model including each constraint metric as a covariate. We assessed predictive power of each model as the area under the curve of the Receiver Operating Characteristic (ROC).

Human pathogenic variants were obtained from ClinVar (Landrum et al 2018). The Ensembl canonical transcripts for SNVs labelled “pathogenic” or “likely pathogenic” were identified using the Ensembl Variant Effect Predictor (v94.5). This left 52,174 pathogenic variants for analysis in human genes with mouse orthologues for which funZ is calculated. Human-mouse orthologues were binned from 1 to 100 based on their funZ percentile, with the least constrained genes in the 1^stt^ bin and the most constrained genes in the 100^th^ bin. To account for differences in gene length, we averaged pathogenic variants in each gene per kb. We fit simple linear regression models to predict the mean number of pathogenic variants per kb as a function of funZ percentile bin for 15,680 mouse and 15,562 human orthologues.

## ACKNOWLEDGMENTS

We would like to acknowledge Hugh Morgan and Luis Santos of MRC Harwell for their feedback and contribution to analysing data from the IMPC.

## AUTHOR CONTRIBUTIONS

C. Lindgren and G. Powell conceived of the presented idea. G. Powell performed the analysis, designed the figures, and wrote the manuscript. All other authors helped support, plan, and supervise the work, and contributed to verifying the analytical methods and producing the final manuscript.

