## Supplementary Information for "Tolerance of nonsynonymous variation is closely correlated between human and mouse orthologues"

**SUPLEMENTARY INFORMATION**

**Functional and non-functional coding annotations**

**Supplementary table 1 --** Coding annotations defined as functional and non-functional from Ensembl’s Variant Effect Predictor (v94.5). One consequence was determined per SNV using the “--pick” argument which prioritises annotations by canonical transcript status.

| **“Functional” annotations** | **“Non-functional” annotations** |
| --- | --- |
| missense_variant | synonymous_variant |
| stop_gained | stop_retained_variant |
| start_lost | start_retained_variant |
| stop_lost | intron_variant |

**Comparison of 1000GP and MGP datasets**

**Supplementary table 2 --** Total canonical transcripts, functional SNVs, and non-functional SNVs in 1000 Genomes Project and Mouse Genomes Project datasets at different MAF cut-offs. To account for the greater proportion of synonymous SNVs in the Mouse Genomes Project dataset, we halved the number of synonymous SNVs in each mouse gene before calculating constraint.

| **Species** | **MAF > threshold** | **N canonical transcripts** | **N synonymous** | **N missense** | **N nonsense** |
| --- | --- | --- | --- | --- | --- |
| Mouse | NA | 19916 | 265111 | 129210 | 831 |
| Human | 0.001 | 18805 | 91770 | 100525 | 1263 |
| Human | 0.0005 | 18811 | 129605 | 155048 | 2194 |
| Human | 0.0001 | 18821 | 342421 | 530701 | 11618 |

**Mutation rates**

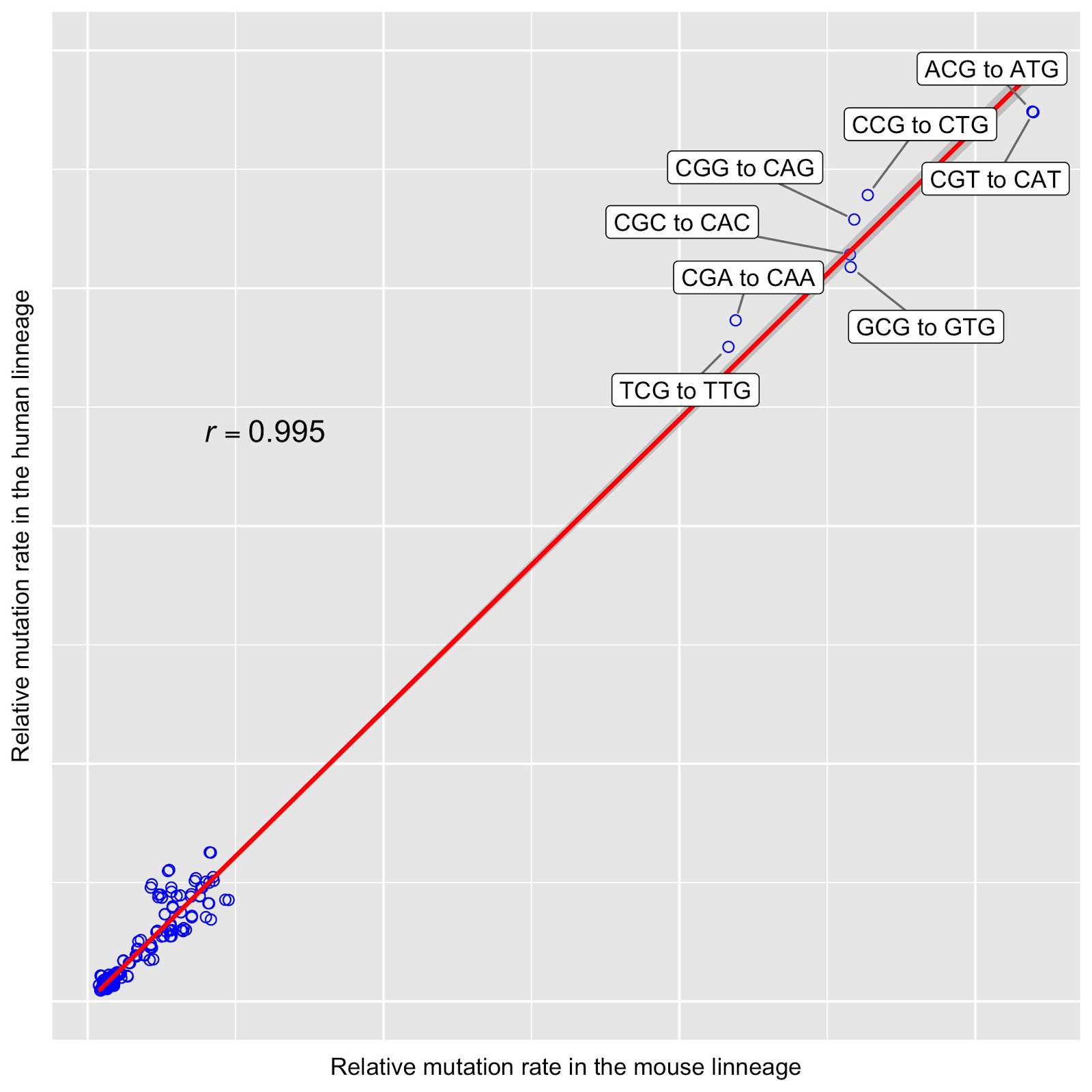

**Supplementary Figure 1 --** Correlation in trinucleotide variation rates between human and mouse lineages. This is largely accounted for by the presence of CpG dinucleotides.

**Predicting the number of synonymous variants**

We built models to predict the number of SNVs in each human and mouse gene assuming no selection pressure by regressing the number of common (MAF > 0.001) synonymous variants in the Mouse Genomes Project and 1000 Genomes Project databases against the genes sequence-specific probability of synonymous mutation, regional mutation rate, and intron mutation rate. Adjusted R2 and mean squared error for the mouse model was 0.748 and 13.7 respectively. Adjusted R2 and mean squared error for the human model was 0.646 and 11.2 respectively. Covariate significance is provided in supplementary table 3.

**Supplementary Table 3 --** Covariate significance for predicting the gene-specific number of synonymous variants (MAF > 0.001) in humans using the 1000 Genomes Project dataset, and mice using the Mouse Genomes Project dataset.

| **Species** | **Covariate** | **Estimate** | **Std. Error** | **P value** |
| --- | --- | --- | --- | --- |
| Human | Sequence-specific probability of synonymous mutation. | 0.261 | 0.002 | <2e-16 |
| Human | Regional mutation rate | 100.086 | 4.794 | <2e-16 |
| Human | Intron mutation rate | 1.614 | 0.133 | <2e-16 |
| Mouse | Sequence-specific probability of synonymous mutation. | 0.474 | 0.002 | <2e-16 |
| Mouse | Regional mutation rate | 148.749 | 7.068 | <2e-16 |
| Mouse | Intron mutation rate | 1.749 | 0.104 | <2e-16 |

**FunZ sensitivity to MAF cutoff**

**Supplementary Table 4 --** Pearson’s product moment correlation coefficients for the correlation between human funZ calculated with MAF > 0.0001, 0.0005, and 0.001 (n = 17,279).

|  | funZ (MAF>0.001) | funZ (MAF>0.0005) | funZ (MAF>0.0001) |
| --- | --- | --- | --- |
| funZ (MAF>0.001) | 1 | 0.98 | 0.87 |
| funZ (MAF>0.0005) | 0.98 | 1 | 0.91 |
| funZ (MAF>0.0001) | 0.87 | 0.91 | 1 |

**Comparison with other measures of intraspecific constraint, and interspecific conservation.**

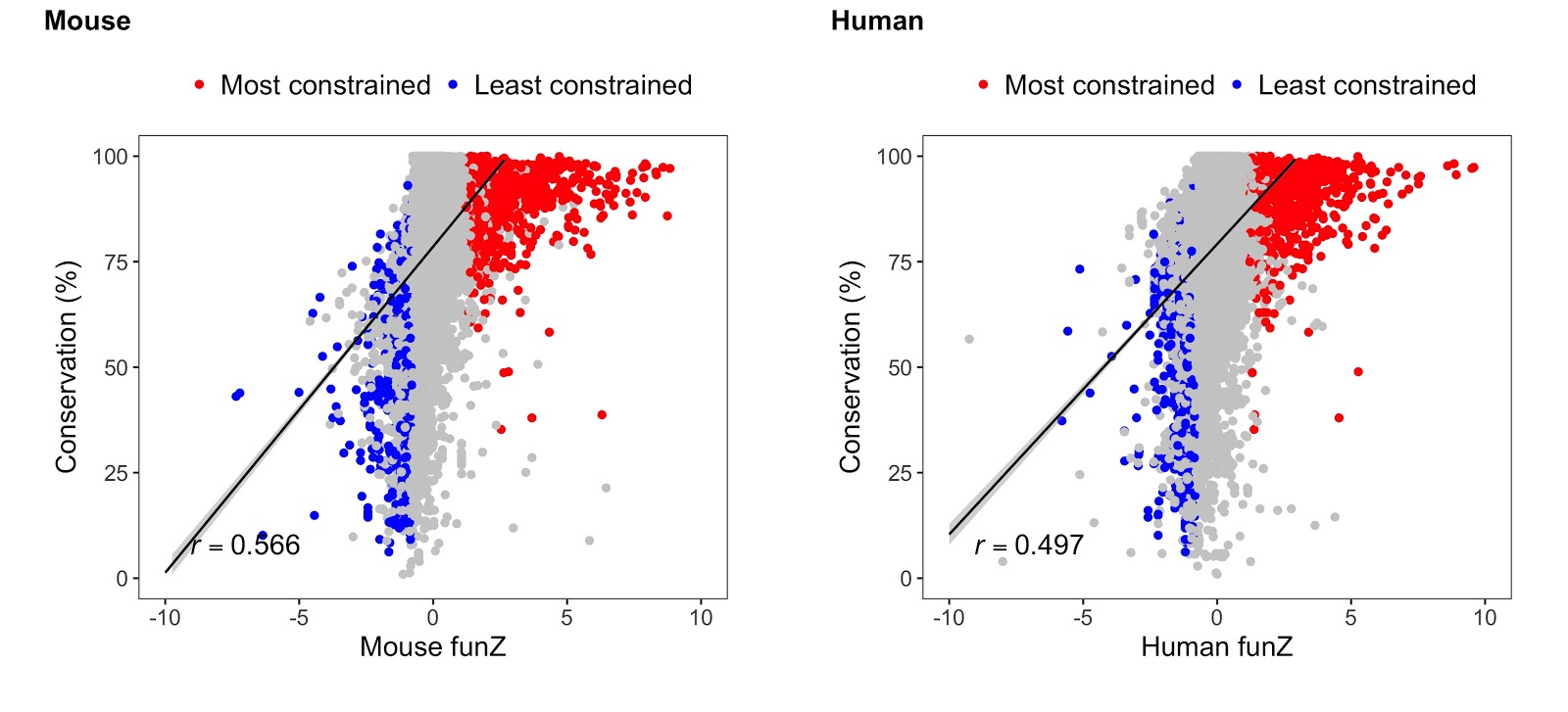

**Supplementary Figure 2 --** Intraspecific constraint is correlated with interspecific conservation. Constraint is measured as funZ, and conservation measured as the percentage of the amino-acid sequence that matches between human and mouse orthologues. Spearman’s Rank correlation coefficients were calculated for 16,270 orthologues. Orthologues annotated in red are amongst the ten percent most constrained (highest funZ) in humans and mice (n = 1,132); orthologues annotated in blue are amongst the ten percent least constrained (lowest funZ) orthologues in humans and mice (n = 751).

**Supplementary Table 5 --** Spearman’s Rank correlation coefficients for the correlation between funZ, missense Z-score, pLI, and RVIS for human genes (n = 14,446).

|  | **funZ** | **RVIS** | **missense Z-score** | **pLI** |
| --- | --- | --- | --- | --- |
| **funZ** | 1 | 0.744 | 0.544 | 0.397 |
| **RVIS** | 0.744 | 1 | 0.444 | -0.382 |
| **missense Z-score** | 0.544 | 0.444 | 1 | 0.622 |
| **pLI** | 0.397 | -0.382 | 0.622 | 1 |

**Knockout phenotype**

**Supplementary Table 6 --** Mann Whitney U test results for the relationships between top-level mouse phenotype terms and knockout constraint.

| Top-level MP term | N knockouts with MP | N knockouts without MP | Mean funZ percentile with MP | Mean funZ percentile without MP | Statistic | P value |
| --- | --- | --- | --- | --- | --- | --- |
| no annotated phenotype | 1339 | 4147 | 0.428 | 0.523 | 2.25e+06 | 2.04e-24 |
| digestive/alimentary phenotype | 63 | 883 | 0.537 | 0.561 | 2.65e+04 | 1.00e+00 |
| endocrine/exocrine gland phenotype | 248 | 270 | 0.506 | 0.568 | 2.92e+04 | 2.70e-01 |
| renal/urinary system phenotype | 147 | 905 | 0.509 | 0.53 | 6.35e+04 | 1.00e+00 |
| integument phenotype | 321 | 4004 | 0.519 | 0.503 | 6.63e+05 | 1.00e+00 |
| cardiovascular system phenotype | 589 | 4092 | 0.525 | 0.501 | 1.26e+06 | 1.00e+00 |
| reproductive system phenotype | 452 | 3768 | 0.499 | 0.492 | 8.64e+05 | 1.00e+00 |
| limbs/digits/tail phenotype | 283 | 4177 | 0.535 | 0.498 | 6.35e+05 | 8.39e-01 |
| embryo phenotype | 253 | 306 | 0.604 | 0.558 | 4.26e+04 | 9.18e-01 |
| homeostasis/metabolism phenotype | 1438 | 3109 | 0.536 | 0.493 | 2.43e+06 | 8.17e-05 |
| immune system phenotype | 677 | 3089 | 0.54 | 0.494 | 1.14e+06 | 3.70e-03 |
| pigmentation phenotype | 73 | 4161 | 0.571 | 0.499 | 1.74e+05 | 7.65e-01 |
| hematopoietic system phenotype | 1045 | 2807 | 0.538 | 0.49 | 1.61e+06 | 8.68e-05 |
| vision/eye phenotype | 472 | 3792 | 0.567 | 0.499 | 1.02e+06 | 3.14e-05 |
| hearing/vestibular/ear phenotype | 175 | 3635 | 0.557 | 0.498 | 3.56e+05 | 1.68e-01 |
| behavior/neurological phenotype | 1387 | 3385 | 0.545 | 0.483 | 2.64e+06 | 4.07e-10 |
| adipose tissue phenotype | 387 | 3089 | 0.567 | 0.49 | 6.89e+05 | 2.10e-05 |
| skeleton phenotype | 784 | 4061 | 0.558 | 0.489 | 1.81e+06 | 1.29e-08 |
| nervous system phenotype | 301 | 3996 | 0.569 | 0.501 | 6.83e+05 | 1.70e-03 |
| growth/size/body region phenotype | 966 | 3887 | 0.582 | 0.48 | 2.26e+06 | 1.32e-21 |
| craniofacial phenotype | 126 | 4162 | 0.621 | 0.497 | 3.27e+05 | 4.59e-05 |
| mortality/aging | 1577 | 3041 | 0.585 | 0.449 | 3.05e+06 | 2.74e-51 |
